# Cell size homeostasis is maintained by a circuitry involving a CDK4-determined target size that programs the cell size-dependent activation of p38

**DOI:** 10.1101/2020.10.14.339556

**Authors:** Ceryl Tan, Miriam B. Ginzberg, Rachel Webster, Seshu Iyengar, Shixuan Liu, John Concannon, Yuan Wang, Douglas S. Auld, Jeremy L. Jenkins, Hannes Rost, Andreas Hilfinger, W. Brent Derry, Nish Patel, Ran Kafri

## Abstract

While molecules that promote the growth of animal cells have been identified, the following question remains: How are growth promoting pathways regulated to specify a characteristic size for each of the different cell types? In 1975, Hartwell and Nurse suggested that in eukaryotes, cell size is determined by size checkpoints – mechanisms that restrict cell cycle progression from cells that are *smaller* than their *target size*. Curiously, such checkpoint mechanisms imply a conceptual distinction between a cell’s *actual* size and cell’s *target* size. In the present study, we materialize this conceptual distinction by describing experimental assays that discriminately quantify a cell’s target size value. With these assays, we show that a cell’s size and target size are distinct phenotypes that are subject to different upstream regulators. While mTORC1 promotes growth in cell size, our data suggests that a cell’s target size value is regulated by other pathways including FGFR3, ROCK2, and CDK4. For example, while rapamycin (an mTORC1 inhibitor) decreases cell size, rapamycin does not change the target size that is required for the G1/S transition. The CDK4/Rb pathway has been previously proposed as a putative regulator of target size. Yet, in lacking experimental means that discriminate perturbations of cell growth from perturbations that reprogram target size, such claims on target size were not validated. To investigate the functions of CDK4 in target size determination, we used genetic and chemical means to ‘dial’ higher and lower levels of CDK4 activity. These measurements identified functions of CDK4 on target size that are distinct from other G1 CDKs. Using *C. elegans*, we further demonstrate that these influences of CDK4 on size determination function *in vivo*. Finally, we propose a model whereby mTORC1, p38, and CDK4 cooperate in a manner that is analogous to the function of a thermostat. While mTORC1 promotes cellular growth as prompted by p38, CDK4 is analogous to the thermostat *dial* that sets the critical target size associated with cell size homeostasis.

## INTRODUCTION

How does signal transduction specify a characteristic cell size for each of the different cell types? Pancreatic acinar cells, for example, are roughly 1.5-fold the size of their neighboring beta cells. Over the past decades, research efforts have identified critical components of cellular growth. Across a wide variety of different tissues, cellular growth is promoted by mTORC1 (Liu and Sabatini, 2020). In addition to mTORC1, other growth promoting signals include, Hippo, cMyc and more (Lloyd, 2013). Yet, despite such progress, a fundamental question stands: How is mTORC1 (or other growth promoting signals) regulated to specify one common size for numerous individual cells within a tissue?

Since the 1970’s, it has become increasingly accepted that the size of eukaryote cells is determined by cell size checkpoints – mechanisms that selectively promote the growth of cells that are smaller than their target size. Initial evidence on size checkpoints was reported by Killander and Zetterberg in 1965 (Killander and Zetterberg, 1965). In that seminal work, mouse fibroblasts were used to show that smaller cells compensate with longer periods of growth in G1, a process that culminates with increased size uniformity in the subpopulation of cells that are undergoing the G1/S transition (Fig. S1B). The concept of a cell size checkpoint was later introduced by showing that a critical mass must be achieved for progression through the cell cycle (Hartwell et al., 1974; Nurse, 1975). In recent years, evidence on size checkpoints have accumulated from multiple independent labs (Darzynkiewicz et al., 1979; Dolznig et al., 2004; Gao and Raff, 1997; Ginzberg et al., 2018; Kafri et al., 2013; Liu et al., 2018; Schmoller et al., 2015; Shields et al., 1978; Tzur et al., 2009). In *S. pombe*, intracellular gradients of Pom1 and Cdr2 were shown to communicate information on cell size and contribute to the control of mitotic entry (Allard et al., 2018; Deng et al., 2014; Gerganova et al., 2019; Martin and Berthelot-Grosjean, 2009; Moseley et al., 2009). However, as mutants in both Pom1 and Cdr2 retain cell size homeostasis, redundant mechanisms may also be in place (Facchetti et al., 2019; Wood and Nurse, 2013). While mechanisms vary, the coordination of cell growth with cell cycle has been shown to be instrumental in the regulation of cell size from bacteria to animal tissues (Conlon and Raff, 2003; Hartwell et al., 1974; Leitao and Kellogg, 2017; Neufeld et al., 1998; Nurse, 1975; Schaechter et al., 1962; Sellam et al., 2019; Willis et al., 2016; Xie and Skotheim, 2020).

Curiously, the functioning of cell size checkpoints implies a conceptual distinction between a cell’s *actual size* and cell’s *target size*. In both yeast and mammalian cells, progression into cell cycle is permissible only when a cell’s *actual size* exceeds a *critical target size value* (Zatulovskiy and Skotheim, 2020). In a previous study, we showed that CDK4 inhibitors promote significant increases in cell size, while still maintaining functional cell size homeostasis (Ginzberg et al., 2018). We hypothesized that CDK4 may have roles in the determination of a cell’s target size. Our hypothesis, however, remained unvalidated. What limited our progress was the absence of adequate experimental criteria for influences on target size. Smaller cells, for example, can result from (1) perturbations that impair mechanisms of cellular growth (e.g. mTORC1 inhibition), or (2) perturbations that dial down the target size associated with cell size homeostasis. How to experimentally discriminate these possibilities presented a challenge.

Recently, we and others have suggested that the p38 MAPK functions to selectively restrict small cells from progressing through the cell cycle (Liu et al., 2018; Sellam et al., 2019). We have also shown that cell size is maintained at its target size value through homeostatic coordination of growth rate and growth duration (Ginzberg et al., 2018). Yet, while recent findings from both ours and other labs have revealed insights, these progress also further emphasize the question: If cells transition into S phase and deactivate p38 only after surpassing a critical size threshold, what then is the mechanism that determines this critical size?

An involvement of the CDK4/Rb pathway in cell size homeostasis has been suggested by several studies. In both yeast and animal cells, cell size sensing was suggested to involve a mechanism that depends on the growth dependent dilution of Rb (Schmoller et al., 2015; Zatulovskiy et al., 2020). Congruently, important roles have been identified for cyclin D1/CDK4 in controlling glucose metabolism and regulating energy balance independently of cell cycle progression (Lee et al., 2014; Lopez-Mejia et al., 2017).

In the present study, we describe two experimental assays that materialize the conceptual distinction between a cell’s *actual* size and cell’s *target* size. To assay changes in target size, we measure: (A) the critical size threshold associated with the G1/S transition and (B) the size threshold associated with the activation of p38 MAPK. With these assays, we show that a cell’s size and target size are distinct phenotypes that are subject to different upstream regulators. While mTORC1 promotes growth in cell size, our data suggests that a cell’s target size value is regulated by other pathways including FGFR3, ROCK2, and CDK4. Specifically, we show that p38 MAPK is selectively activated in cells that are smaller than a critical target size. We further show that the critical cell size threshold that discriminates active and inactive forms of p38 is ‘dialed’ to higher or lower values by changes in CDK4 activity. By contrast, rapamycin (an mTORC1 inhibitor) increases the proportion of small, p38 active cells, but does not change the target size associated with p38 activity or the G1/S transition. Finally, we propose a model whereby – just as a thermostat maintains room temperature – CDK4 and p38 function in a circuitry that maintains cells within a programmed size range (Fig. S1A). Specifically, we suggest that while p38 is selectively activated in cells that are smaller than their target size, CDK4 functions to determine the target size associated with p38 activation and entry into cell cycle. To the best of our knowledge, this is the first report outlining experimental assays that has materialized target size from a concept to a measurable phenotype that is separate from cell size.

## RESULTS

### Cell size homeostasis is maintained by a reciprocal coordination of growth rate and growth duration

In proliferating cells, size checkpoints function to coordinate rates of cell division with rates of biosynthetic activity in order to attain the specified target size (Cadart et al., 2018; Ginzberg et al., 2018; Leitao and Kellogg, 2017). These constraints result in a negative correlation between a cell’s birth size and the length of its G1 phase (Liu et al., 2018; Varsano et al., 2017; Xie and Skotheim, 2020). This reciprocal coordination is observed in unperturbed conditions and also persists across a wide variety of chemical and genetic perturbations (Ginzberg et al., 2018; Liu et al., 2018). To characterize cell size checkpoints, we treated RPE1 cells with increasing doses of rapamycin (mTORC1 inhibitor). To measure cell size, we labeled cells with SE-A647 (Fig. S2) as previously described (Ginzberg et al., 2018; Kafri et al., 2013; Liu et al., 2018), and quantified fluorescence intensity from the nuclear or cytoplasmic boundaries (Methods). Consistent with (Ginzberg et al., 2018), rapamycin promotes dose-dependent decreases in rates of biosynthesis (growth rate) that are accompanied with reciprocally proportional changes in G1 length, resulting in only modest changes in cell size (Fig. 1A). These coordinate influences of rapamycin on growth rate and G1 length are consistent with a cell size checkpoint.

**Figure 1.**
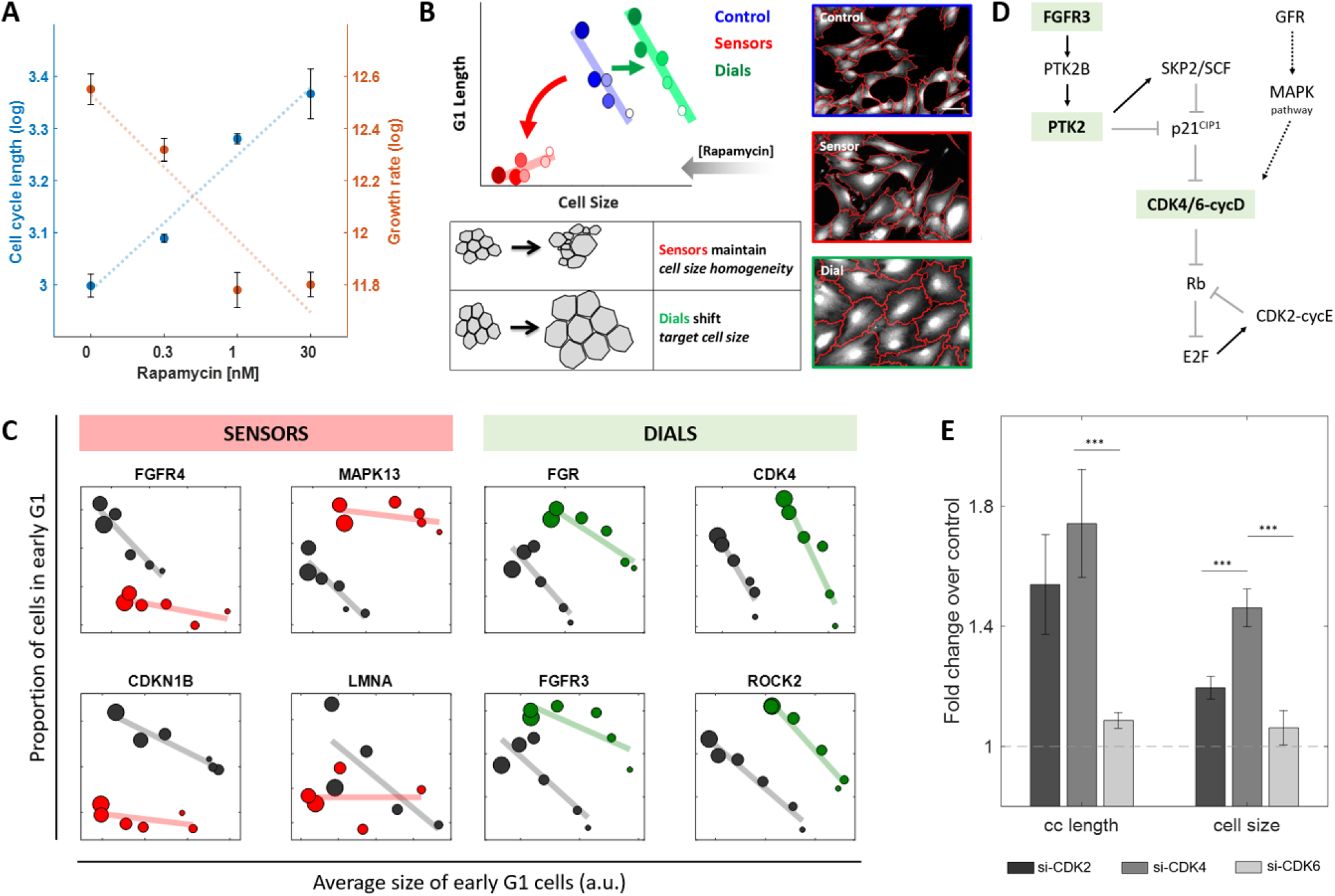
Growth rate has a quantitative, reciprocal influence on growth duration. **(A)** Growth rate (*v*, protein synthesis per unit time) and growth duration (*v*, cell cycle length) for RPE1 cells treated with increasing doses of rapamycin. Both growth rate and growth duration values are represented on log scale to emphasize the conservation of cell size, log(*S*) = *log*(*τ* × *v*) = log(*τ*) + log (*v*). Data are from two distinct samples and presented as median ± SEM. **(B)** Unsynchronized RPE1 cells were treated with increasing concentrations of rapamycin for 24 hr. Each data point (circle) corresponds to a different concentration of rapamycin, quantified for the average size of early G1 cells and the proportion of cells in G1 (Methods). Perturbations that disrupt the coordination between cell size and G1 length were classified as “sensor” perturbations, while compounds that shifted the coordination to higher or lower setpoint values were categorized as “dials”. Phenotypically, disrupting sensors result in a loss of cell size homogeneity, while perturbing dials result in homogeneously smaller or larger cell sizes across the population. Representative widefield fluorescence images (SE-A647 stain) of RPE1 cells are shown at the same scale, scale bar = 50μm. Red lines delineate computationally segmented cell boundaries. **(C)** RPE1 cells transfected with siRNA targeting representative sensors (red) and dials (green) as well as si-CONTROL (black) and their influence on the rapamycin-induced negative correlation between cell size and G1 length. **(D)** Targets that resulted in a dial phenotype (highlighted in green) form a pathway that is parallel and functionally redundant with CDK4/6-cycD. **(e)** Fold-change in cell cycle (cc) length and cell size for RPE1 cells transfected with si-CDK2, si-CDK4, or si-CDK6, compared to si-CONTROL. Data are from three independent experiments and presented as median ± SEM. (*\*\*\*p-value<0.05* from two-sample t-test).

### Experimental assay for dials and sensors of cell size

To investigate mechanisms of size determination, we systematically characterized perturbations that were previously suggested to promote changes in cell size homeostasis (Liu et al., 2018). We considered two general possibilities: (1) a departure from cell size homeostasis can result from perturbations that disrupt the proper functioning of cell size checkpoints. In such cases, cells will transition into S phase in a manner that is independent of their size, resulting in increased cell size variance (Fig. 1B) (Liu et al., 2018). Alternatively, (2) deviations from cell size homeostasis can also result if properly functioning checkpoints are reprogrammed for higher or lower target size values (Fig. 1B). To discriminate mechanisms that enable cell size checkpoints from mechanisms that program the target size associated with these checkpoints, we screened for perturbations that affect the linear dependency of cell size with G1 length (Fig. 1B). For simplicity, we will refer to these categorical distinctions with the terms: *dials* and *sensors* of cell size. Specifically, we define *sensors* as perturbations that weaken the dependency of size on G1 length. By contrast, *dials* were classified as perturbations that affect cell size but do not weaken the coordination of size and G1 length (Fig. 1B). Compounds that produced a statistically significant dial or sensor phenotype were selected for further validation using siRNA (Fig. 1C).

### A chemical screen identifies CDK4 as a putative dial of cell size

One of the top sensors identified from this assay was MAPK13 (p38*δ*), which is in line with our previous findings (Liu et al., 2018), as well as a recent study in *C. albicans* (Sellam et al., 2019). As top candidates for the dial phenotype, we obtained CDK4, FGFR3, and PTK2. In previous studies, FGFR3 and PTK2 have been implicated as a branch of signaling that functions in parallel, and is partially redundant with, the CDK4/6-Rb pathway (Fig. 1D) (Zhang et al., 2019). The possibility that the CDK4/6-Rb pathway functions in target size determination and cell size homeostasis is further supported by previous literature (Meyer, 2000; Schmoller et al., 2015; Zatulovskiy et al., 2020) and work from our own lab (Ginzberg et al., 2018). Curiously, the CDK4/6-Rb-E2F pathway has recently been implicated with promoting metabolic activity in a manner that is independent of its roles in cell cycle (Franco et al., 2016; Lee et al., 2014; Wang et al., 2006).

### CDK4 activity determines the quantitative relationship between cell size and G1 length

Canonically, CDK4 is a cyclin-dependent kinase (CDK) that functions upstream to CDK2 and is partially redundant in function with its isoform, CDK6. A central role of the G1 CDKs is to promote the phosphorylation of the retinoblastoma protein (Rb), thus allowing passage through G1/S (Malumbres and Barbacid, 2009). Since CDK2, CDK4, and CDK6 have overlapping functions, we sought to characterize each for their influence on G1 length and cell size. While knockdown (KD) of CDK2 promoted a ~60% increase in the duration of G1, the influence of CDK2 KD on cell size was only ~20% (Fig. 1E, Fig. S3C). Naively, cells that grow for longer periods of time should become proportionally larger in size. The relatively modest influence of CDK2 KD on cell size, as compared to its influence on G1 length alludes to a coordinate reduction in growth rate (Ginzberg et al., 2018). As with the knockdown of CDK2, a significant lengthening of G1 (~70%) was also observed with siRNA against CDK4. However, as compared to CDK2, depletion of CDK4 promoted significantly larger increases in cell size (Fig. 1E, Fig. S3C). Last, KD of CDK6 resulted in a relatively modest influence (<10%) on both G1 length and cell size. It is also worth noting that although knocking down the levels of these kinases almost doubles cell cycle length, cells remain in exponential proliferation for up to 96-hours post-transfection (Fig. S3A-B). Overall, these results suggest that the different G1 CDKs serve divergent roles in cell size regulation, with CDK4 potentially playing a distinct role in target size regulation.

If CDK4 functions to quantitatively specify the target size associated with the G1/S transition, decreasing CDK4 activity should promote gradual shifts in the correlation of cell size and G1 length. To reduce CDK4 activity, we exposed cells to palbociclib, a chemical inhibitor of CDK4/6. At concentrations that exceed 1μM, palbociclib promotes cell cycle arrest (Llanos et al., 2019). For our experiments, we carefully optimized a palbociclib concentration range that decreases CDK4 activity but does not result in cell cycle arrest (Fig. S4). Indeed, increasing concentrations of palbociclib promoted a dose-dependent shift in the correlation of cell size with G1 length (Fig. 2A). Similar shifts were observed on primary human fibroblasts, as well as using a different CDK4/6 inhibitor, abemaciclib (Fig. S5). Given the modest increase in cell size observed with CDK6 knockdown (Fig. 1E, Fig. S3C), we speculate that the up to 60% change in cell size observed with palbociclib is mediated by CDK4. Since reductions in CDK4 activity resulted in larger cells, we asked whether increased CDK4 activity would promote the opposite effect, causing downward shifts in target size. To increase the activity of CDK4, we generated a stable RPE1 cell line with doxycycline-inducible overexpression of cyclin D1 (dox-cycD1), which forms an active complex with CDK4/6 (Ginzberg et al., 2018). Consistent with our expectations, increasing concentrations of doxycycline resulted in smaller cell sizes, in a dose-dependent manner, while still maintaining a tight correlation of cell size and G1 length (Fig. 2A).

**Figure 2.**
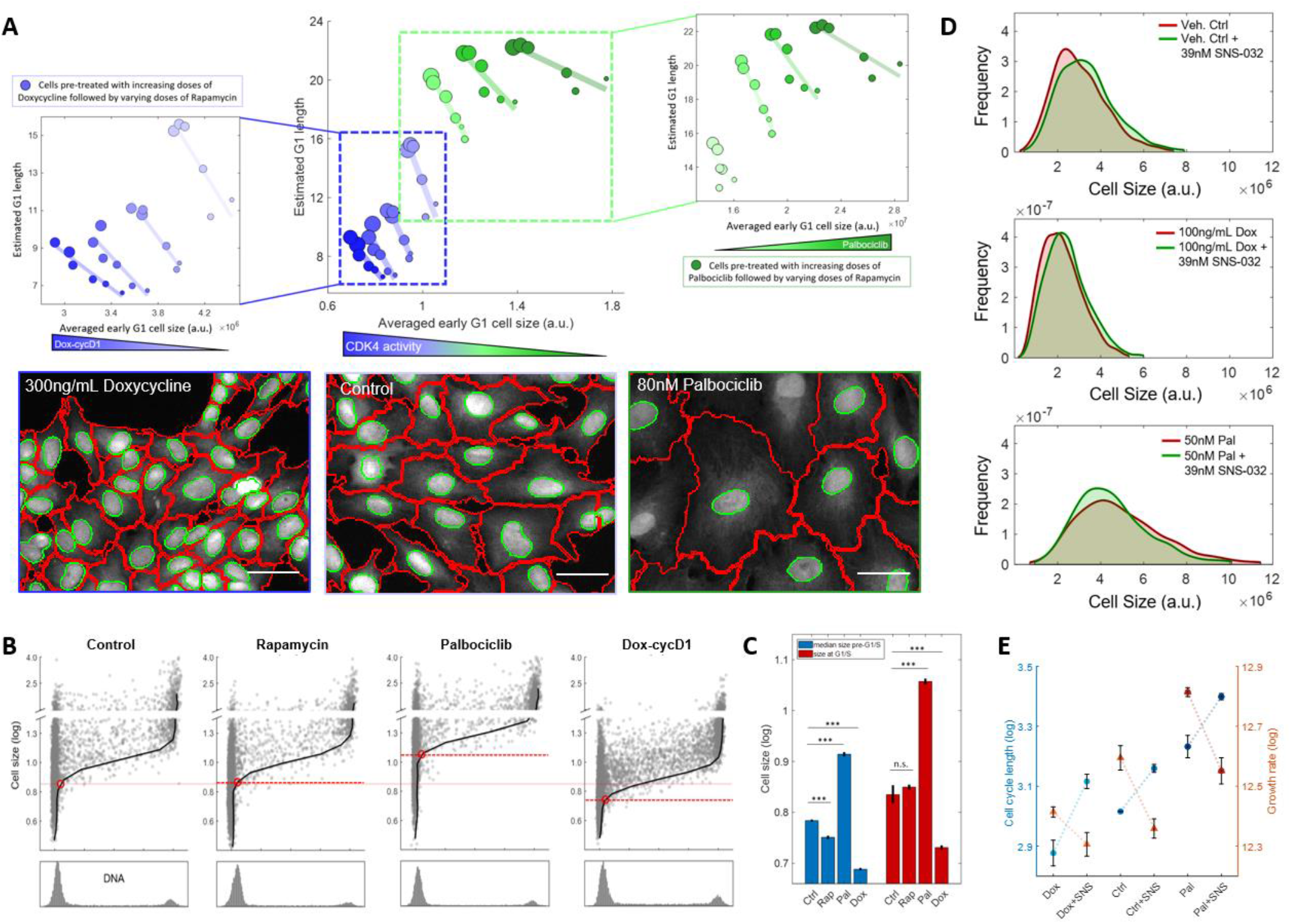
CDK4 activity influences the association between cell size and G1 length. **(A)** RPE1 cells were pre-treated with 0, 150, 300, or 400ng/mL of dox-cycD1 (blue) or 0, 10, 40, or 80nM of palbociclib (green) for 24 hrs, followed by varying concentrations of rapamycin for another 24 hrs (Methods). Representative widefield fluorescence images (SE-A647 stain) are shown at the same scale, scale bar = 25μm. **(B)** To highlight the cell size criterion associated with the G1/S transition, red lines demarcate the size at which the population transitions from G1 to S (DAPI=2.1). Progression through the G1/S transition was observed only in cells whose size (nuclear SE-A647) exceeded a given threshold (red lines). This threshold was unchanged by mTORC1 inhibition (30nM Rap) but was shifted to higher (25nM Pal) or lower (100ng/mL Dox) values with CDK4 perturbations. Solid black lines indicate the DNA level that associates with the 95th percentile larger cells per DAPI value (Methods). **(C)** Median sizes prior to G1/S and cell size at the G1/S transition are plotted for the various drug treatments. Error bars indicate 95% CI, \*\*\**p-value* <0.001 (two-sample t-test). Ctrl: n=9, Rap: n=3, Pal: n=3, Dox: n=3. **(D)** RPE1 cells were pretreated with compounds to increase (dox) or decrease (pal) CDK4 activity for 24 hrs, after which cells were incubated with 39nM SNS-032. Cell size distributions are presented for the different combinations of drug treatment at 40 hrs post-SNS treatment. **(E)** Cell cycle length and growth rate are presented on log scale for the different combinations of activities in CDK4 (dox, ctrl, pal) and CDK2 (39nM SNS-032). Data are from three distinct samples and presented as median ± SEM.

In proliferating cells, progression into S phase is regulated by a target size threshold (Dolznig et al., 2004; Gao and Raff, 1997; Ginzberg et al., 2018; Kafri et al., 2013; Liu et al., 2018; Schmoller et al., 2015; Shields et al., 1978; Tzur et al., 2009). To investigate whether CDK4 determines this threshold, we performed single cell measurements of size and cell cycle stage in individual cells. In unperturbed conditions, RPE1 cells transition into S phase at a clearly delineated target size (Fig. 2B). While both rapamycin and the activation of CDK4 (dox-cycD1) promoted downward shifts in cell size, it is only the latter that further decreased the target size associated with the G1/S transition (Fig. 2B-C). Conversely, reducing CDK4 activity with palbociclib promoted the opposite effect, with cells transiting into S phase at a higher target size value (Fig. 2B-C). Collectively, these results demonstrate that fine-tuning CDK4 activity can quantitatively reprogram the relationship between cell size and G1 length, such that target size can be dialed up or down by modulating CDK4 activity. These results support the conceptual distinction between a cell’s *size* and its *target size*, suggesting that these are two separately regulated phenotypes.

### CDK4 specifies the proportionality constant relating cell cycle length with rates of biosynthesis

Our data suggest that CDK2 and CDK4 play different roles in cell size homeostasis (Fig. 1E). While both kinases regulate G1 length, only CDK4 promotes significant changes in the target size associated with the G1/S transition. To investigate whether CDK2 and CDK4 bear synergistic effects on cell size homeostasis, we tested the influence of CDK2 inhibition in cells with increased (dox-cycD1) or reduced (palbociclib) CDK4 activity. Inhibiting both CDK2 and CDK4 activities resulted in a longer G1 phase than either alone (Fig. S6). Yet, while CDK2 inhibitors further increased the cell cycle length of cells already subject to palbociclib treatment by over 60%, the influence of CDK2 inhibitors on the size of palbociclib-treated cells was only 6% (Fig. 2D, Fig. S6). This relatively small influence on cell size is because the *longer periods* of growth (longer cell cycles) caused by the inhibition of CDK2 were mitigated by *slower rates* of biosynthesis (Fig. 2E). Strikingly, these mutual influences of CDK2 on growth rate and growth duration were proportionally coordinated to result in smaller or larger target size values, depending on levels of CDK4 activity. These observations are not restricted to CDK2 inhibition but extend to an array of inhibitors against growth rate and growth duration (Fig. S6). Across multiple perturbations, palbociclib and dox-pretreated cells maintained a constant target size that was ~25% larger or smaller than control, respectively (Fig. 2D, Fig. S6).

### To maintain cell size homeostasis, CDK4 and p38 function in a circuitry that is analogous to the function of a thermostat

Our experiments suggest a conceptual distinction between a cell’s size and a cell’s target size. To further investigate cells that are smaller than their target size, we performed measurements on cells recovering from torin2 treatment. In RPE1 cells, torin2 promotes a 20% decrease in cell size. Following torin2 wash out, mTORC1 resumes activity almost immediately (Liu et al., 2018). By contrast, the buildup of cell mass and resumption of target size is a slow process that proceeds gradually, over a period of close to 20 hours (Liu et al., 2018). These dynamics provide a significant time window for measurements on cells that have functioning mTORC1 but are still smaller than their target size. If CDK4 reprograms target size, we expect cells released from torin2 inhibition to recover to different target size values depending on the levels of CDK4 activity. Consistent with our understanding, RPE1 cells with activated or repressed CDK4 activity recovered to modified target size values that were respectively smaller or larger than control and stabilized at this new size for at least 2 cell cycles (Fig. 3A).

**Figure 3.**
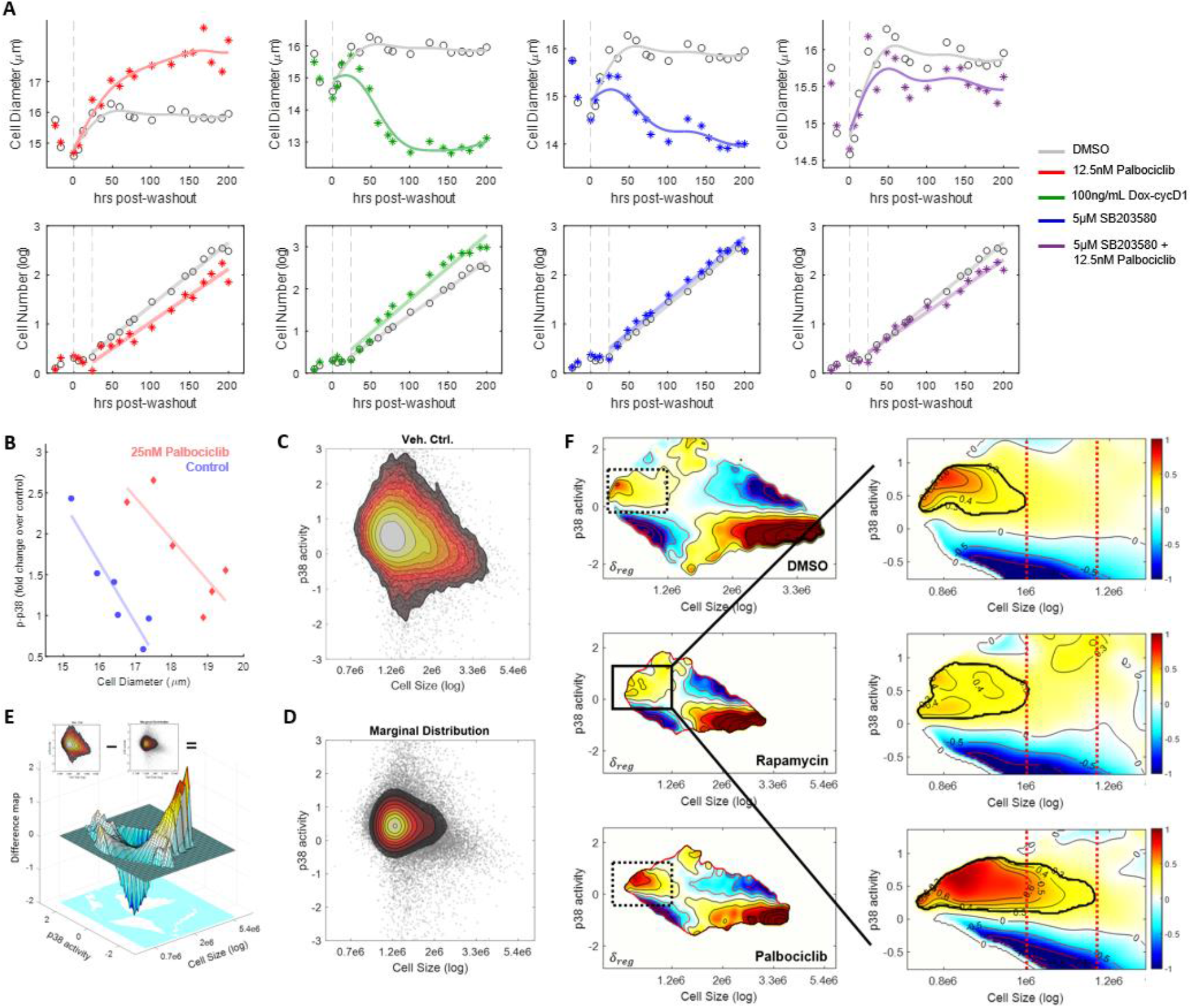
CDK4 activity dials the size threshold at which p38 is activated. **(A)** Cell size (diameter) measurements were performed on RPE1 cells during and after the release from a 20 hr torin2 treatment. Measurements were performed on cells with reduced (red, palbociclib) or increased (green, dox-cycD1) CDK4 activity, reduced p38 activity (blue, SB203580), or with both CDK4 and p38 activity reduced (purple). Vehicle control (0.1% v/v DMSO) is presented in gray. The perturbations influence cell proliferation, but growth is still maintained at an exponential state. **(B)** Cells were pre-treated with 25nM palbociclib or vehicle control for 24 hr prior to a 20 hr treatment of torin2. After washout of torin2, cells were allowed to recover in conditioned media containing either palbociclib or vehicle control. Cell size (diameter) and p38 activity (normalized to washout control for every timepoint) were measured for cells at different stages of recovery. **(C)** Scatter plot overlaid with a contour of the joint probability density distribution of cell size (nuclear SE-A647) and p38 activity (p38-KTR) for control and palbociclib-treated cells. **(D)** The product of the marginal probability distributions, *f*_*ind*_ = *f*(*size*) × *f*(*p38KTR*) **(E)** The log ratio of the two density maps from (C) and (D) creates the difference map, 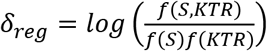. **(F)** *δ*_*reg*_ calculated from single cell measurements on RPE1 cells that were treated with DMSO (control), rapamycin or palbociclib. The blue-red colorscale represents combinations of p38-KTR and cell size that are overrepresented (red) or underrepresented (blue) in the single cell measurements, as compared to the null hypothesis whereby size and p38 are independent. Inset highlights the smaller cells in the population. Red lines illustrate the size at which overrepresented p38 activity cease (bolded black contour line 0.3).

Previously, we showed that p38γ/δ are selectively activated in cells that are smaller than their target size (Liu et al., 2018). To monitor the dynamics of p38 activity throughout recovery from torin2, we collected whole cell lysates at different time points and probed for p38 activity (Fig. S7). Consistent with our previous report, p38 maintained levels of activity that inversely mirrored the slow and gradual recovery in cell size (Liu et al., 2018). After torin2 washout, and despite the resumption in mTORC1 activity, p38 maintained an active state so long as cells were smaller than their CDK4-determined target size. These dynamics resulted in an inverse correlation between cell size and p38 activity (Fig. 3B). When measurements were performed on cells with reduced CDK4 activity, the inverse relationship between size and p38 activity was maintained but shifted to higher cell size values. In control cells, p38 remained active throughout size recovery and only resumed an inactive state when a target size of ~17μm (cell diameter) has been attained. In contrast, palbociclib-treated cells with the exact same size maintained active p38 and was still increasing in size (Fig. 3B).

We next asked whether CDK4 can also reprogram the cell size dependency of p38 activation in unperturbed cells. We utilized a p38 Kinase Translocation Reporter (p38-KTR) that quantitatively reports p38 activity in single cells (Regot et al., 2014). With this system, we performed single cell measurements of size and p38 activity to calculate the joint probability density function, *f*(*S*, *KTR*) (Fig. 3C). One weakness of the p38-KTR system, as described by (Regot et al., 2014), is that the p38-KTR signal intensity that discriminates active and inactive forms of p38 varies across individual cells in a given population. In previous studies, time lapse imaging was used to score each individual cell for its maximal and minimal p38-KTR intensity values (Regot et al., 2014). This can significantly limit cell count and, therefore, the detection of rare events such as that of a small sub-fraction of cells that are both close to the G1/S transition and smaller than their target size. Further, we assumed that measured p38-KTR values may be a function of size dependent mechanisms (*s*), but also size independent mechanisms (*ns*), and noise (*r*), i.e. *p38KTR* = *Ψ*(*s*, *ns*, *r*) (Eq. 1, Box 1). To isolate the specific dependency of p38 activity on cell size, we normalized *f*(*S*, *KTR*) to the product of the marginal probabilities (Fig. 3d): 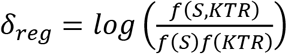 (Eq. 2, Box 1). Effectively, *δ*_*reg*_ quantifies whether cells with any combination of size and p38-KTR are over- or underrepresented in our data (Fig. 3E). By measuring δ_*reg*_ in cells that are subject to different drug treatments, we can investigate how different perturbations affect the target size threshold associated with p38 activation (Fig. 3F).

Consistent with our previous findings, this analysis demonstrates a significant association of p38 activity with small cells (Fig. 3F). Next, we repeated single cell measurements of size and p38-KTR on cells with perturbed activity in mTORC1 or CDK4. Strikingly, the dependency of p38 activity on size was altered by palbociclib but was almost unaffected by rapamycin (Fig. 3F). While rapamycin changed cell size, it did not affect the size threshold associated with the activation of p38. On the other hand, the cell size threshold that discriminates active and inactive forms of p38 was reprogrammed to larger cell size values when CDK4 activity was reduced. Together, these results further discriminate cell size and target size as separate phenotypes, the latter of which can be dialed by CDK4 activity.

If the coordination of a cell’s size and CDK4-determined target size is mediated by p38, then treatment with SB203580 (p38 inhibitor) should prevent CDK4 from reprogramming target size. To test this, we relied on the torin2 recovery protocol. In Fig. 3A, we show that cells relieved from torin2 treatment recover to different target size values, depending on levels of CDK4 activity. Indeed, palbociclib-treated RPE1 cells recovering from torin2-mediated size changes were unable to recover to their new target size in the presence of p38 inhibition (Fig. 3A). Interestingly, while the inhibition of p38 has been shown to increase cyclin D1 expression (Casanovas et al., 2000; Thoms et al., 2007), which may be negating the effects of palbociclib, we show that CDK4 activity remains significantly reduced when co-treated with SB203580 as proliferation rates continuously lagged behind that of control (Fig. 3A). This may be significant because SB203580 by itself increases proliferation rate (Liu et al., 2018).

### CDK4 activity changes cell size in animal tissues

Our results suggest that CDK4 functions to determine target size in animal cells. In *C. elegans*, CDK4 and cyclin D1 are highly conserved as *cdk-4* and *cyd-1*, respectively. To determine whether our findings are relevant *in vivo,* we examined the influence of CDK4 on cell size in the worm. During development, *C. elegans* body growth is driven by both cell size and cell number. In adulthood, somatic cells are terminally differentiated, and therefore any further growth is directly proportional to an increase in cell size (Flemming et al., 2000). Nuclear and nucleolar sizes scale with cell size and play important roles in cell size homeostasis in the worm; therefore, we use the size of these organelles as a proxy for cell size (Altun and Hall, 2002; Weber and Brangwynne, 2015).

Expression of both *cdk-4* and *cyd-1* is observed in a number of tissue types through larval development but is enriched in hypodermal seam cells (Park and Krause, 1999). We therefore knocked down *cdk-4* and *cyd-1* in worms expressing GFP under a seam cell-specific promoter. This allowed for the simultaneous visualization of both nucleoli and nuclei. Multi-generational KD by RNA interference (RNAi) resulted in adult worms that remained viable and fertile, but had enlarged seam cell nucleoli and nuclei (Fig. 4A-B). The variance of nucleolar and nuclear sizes both within and between worms increased significantly compared to controls, however this may be attributable to variations in RNAi KD efficiency. Curiously, the ratio of nucleolar to nuclear areas remained comparable to controls despite the overall increase, confirming that these changes in size are uniform (Fig. 4C). Although there are slight variations in the total number of seam nuclei upon knockdown of either *cdk-4* or *cyd-1*, alluding to their canonical role in cell cycle progression, the average number of nuclei under each RNAi condition were held constant compared to controls (Fig. S8A). Interestingly, body sizes of both *cdk-4* and *cyd-1* were smaller than control animals (Fig. S8B-C). While we did not anticipate this observation, it is in line with the results of a previous study in *Drosophila* (Meyer, 2000).

**Figure 4.**
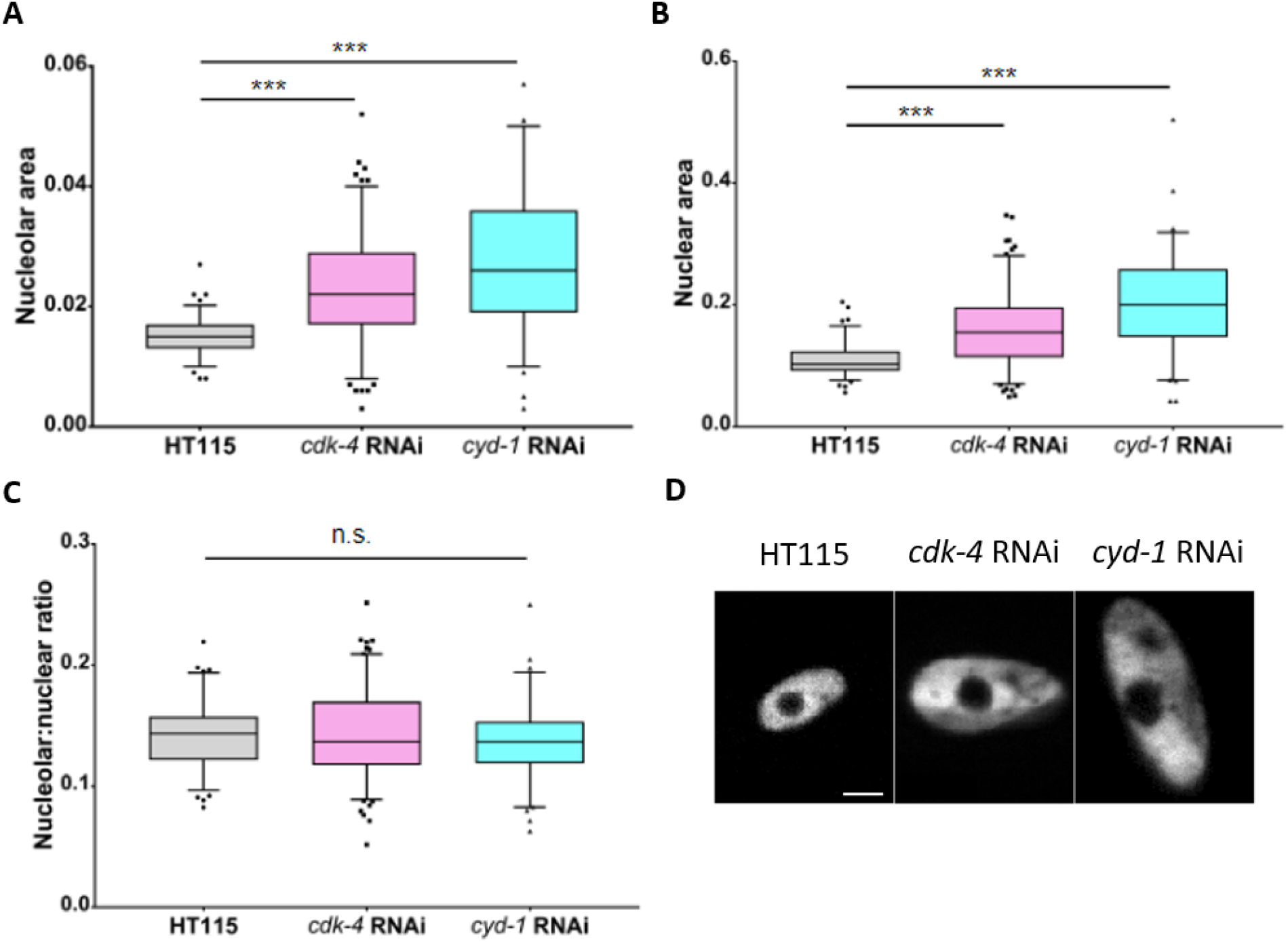
CDK4 activity changes cell size in animal tissues. *C. elegans* Hyp 7 seam cell nucleolar **(A)** and nuclear **(B)** cross-sectional areas increase with knock down of *cdk-4* or *cyd-1* by RNAi feeding compared to empty vector (HT115) controls, measured one day post-adulthood (1DPA). **(C)** The ratio between nucleolar area and its corresponding nuclear area remains constant despite size changes. **(D)** Representative images of labelled seam cell nuclei from the different conditions. Hollowed out regions are nucleoli. Scale bar = 8μm. HT115 RNAi n=6 worms, *cdk-4* RNAi n = 9, *cyd-1* RNAi n = 5. ***p<0.0001, unpaired t-test.

## DISCUSSION

It is well established that mTORC1 promotes cell growth. In (Liu et al., 2018), we showed that p38 is selectively activated in cells that are smaller than their target size. While our work implicated p38 to be involved in cell size sensing, we do not yet have definitive evidence that it indeed is the sensor. Nevertheless, in the present study, we show that the target size referenced to by p38 is programmed by levels of CDK4 activity. Altogether, our results suggest a narrative whereby mTORC1, p38, and CDK4 function in a circuitry that is analogous to a thermostat (Fig. S1A).

Based on our results, we suggest a conceptual distinction between a cell’s size and its target size. This contrast between cell size and target size is illustrated by comparing the following two results: 1) While rapamycin resulted in a smaller cell size, rapamycin did not influence the target size associated with the G1/S transition (Fig. 2B-C) nor the threshold that discriminates active and inactive forms of p38 (Fig. 3B,F). The selective influence of rapamycin on cell size, but not target size, suggests that the latter may be separate from mTORC1 activity. However, such a conclusion demands future studies and genetic analyses of mTORC1 pathway components. 2) We presented two separate populations of cells that are indistinguishable in average cell size (~17μm) but have different target size values (Fig. 3B). At this size, the population with normal CDK4 activity has resumed baseline levels of p38 activity, while the population with reduced CDK4 activity exhibited activated p38 and was still increasing in size. While the first example illustrates how cells of different sizes can still achieve the same target size, the latter exposes cells with equal sizes but different target size.

Another interesting observation from our study is the distinct roles of CDK2 and CDK4 on cell size homeostasis. Since these CDK’s function in a linear pathway, one may expect similar phenotypes. Nonetheless, the influence of CDK4 (but not CDK2) on biosynthetic metabolism was revealed in previous studies. For example, it was reported that CDK4 control glucose metabolism independently of cell cycle progression (Helman et al., 2016; Lee et al., 2014). Further, CDK4 was reported to directly influence growth by phosphorylating AMPKα2 (Lopez-Mejia et al., 2017). The role of the CDK4/cyclin D-Rb axis on cell size regulation has been purported by studies from our lab and others (Ginzberg et al., 2018; Schmoller et al., 2015; Zatulovskiy et al., 2020). While the mechanisms by which CDK4 activity sets target size remains to be investigated, there is increasing evidence that inhibition of CDK4 activity results in increased biosynthetic capacity, such as mitochondrial size and activity, presumably through its feedback on mTORC1 signaling (Cretella et al., 2018; Franco et al., 2016; Haines et al., 2018; Jansen et al., 2017; Romero-Pozuelo et al., 2020).

To conclude, we have shown, to the best of our knowledge, the first evidence for target cell size regulation in animal cells. An increase or decrease in CDK4 activity results in smaller or larger target sizes, respectively, while still maintaining mechanisms of homeostatic size regulation. These results also support the model of cell autonomous size sensing in animal cells, wherein cells can actively quantify their size, compare it against a given setpoint, and adjust accordingly.

## ACKNOWLEDGEMENTS

We thank all the members of the R.K. laboratories for helpful discussions and support. This work was supported by grants to R.K. from the Canadian Institutes of Health Research (343437) and the Natural Sciences and Engineering Research Council of Canada (RGPIN-2015-05805). We also thank Patricia and Alexander Younger and the Younger Foundation for their generous donation to support this research. C.T. was supported by a University of Toronto Open Fellowship. M.B.G. was supported by a postdoctoral fellowship from the Research Training Center at The Hospital for Sick Children.

## AUTHOR CONTRIBUTIONS

C.T. and R.K. conceived the study. C.T., S.L., J.C., Y.W., D.A., and J.J. performed and analyzed the high content chemical screen. C.T., M.B.G., R.W., W.B.D. and N.P. designed and performed the experiments. C.T., M.B.G., H.R., S.I., A.H., and R.K. supported in the analytical methodology development. C.T., N.P., R.W., and R.K. wrote the manuscript with input from all authors.

## METHODS

### Materials

This study did not generate new unique reagents. Palbociclib, Rapamycin, and SB203580 were purchased from Selleckchem. Lentiviral expression vector encoding the p38 MAPK Kinase Translocation Reporter (KTR) was a kind gift from Markus Covert (Addgene plasmid no. 59155). ON-TARGETplus SMARTpool siRNAs for the genes of interest as well non-targeting negative control siRNAs were obtained from Dharmacon (Lafayette, CO). To slow growth rate, the following drug treatments were used: cycloheximide (Sigma C4859), Torin-2 (Tocris 4248), rapamycin (CalBiochem 553211). To slow the cell cycle, the following drug treatments were used: BN82002 (Calbiochem 217691), SNS-032 (Selleckchem S1145), PHA848125 (Selleckchem S2751), Cdk2 Inhibitor III (Calbiochem 238803), Dinaciclib (Selleckchem S2768). All primary (p-p38 Thr180/Tyr182: 4511, p-Rb S795: 9301, p-4EBP1 Thr37/46: 2855) and secondary antibodies used for Western Blotting were purchased from Cell Signaling Technology (Beverly, MA).

### Cell culture

Retinal pigmented epithelial (RPE1, ATCC, RRID:CVCL 4388) cell lines stably expressing the degron of Geminin fused to Azami Green were cultured in DMEM medium (Life Technologies) supplemented with 10% Fetal Bovine Serum (FBS, Wisent, Montreal, QC) at 37°C in a humidified atmosphere with 5% CO_2_. Measurements were made when cells were 50–75% confluent, to avoid the effects of sparse or dense culture on cell growth and proliferation. Detailed methods on the generation of doxycycline-inducible overexpression of cyclin D1 in RPE1 cells can be found in Ginzberg et al(Ginzberg et al., 2018).

### Fixation, staining and imaging

Cells were fixed in 4% paraformaldehyde (Electron Microscopy Sciences, Hatfield, PA) for 10 min, followed by permeabilization in cold methanol at −20°C for 5 min. Cells were stained with 0.4 ug/mL Alexa Fluor 647 carboxylic acid, succinimidyl ester (SE-A647, Invitrogen A-20006), for 2 hours to nonspecifically label total protein. DNA was stained with 1 μg/mL DAPI (Sigma D8417) for 10 min. The cells were imaged using the Operetta High-Content Imaging System (PerkinElmer, Woodbridge, ON) at 20X magnification.

### Chemical screen

The chemical screen described in this text is a follow-up of one described in Liu et al.(Liu et al., 2017), to which we refer readers for more details. Briefly, the primary screen identified compounds that resulted in disproportional changes in either cell size or G1 length (*off-axis compounds*)^12^. We further investigated these compounds using internal Novartis libraries sourced from the publicly known subset of compounds in the Mechanism-of-Action (MOA) Box(Canham et al., 2020). The screen was performed in 384-well μclear microplates (Greiner Bio-one, Monroe, NC). On day 1, 1000 cells per well were seeded in 20 μl medium. On day 2, compound was added to a final concentration of 1 μM, 3 μM or 10 μM from a 2 mM or 10 mM stock solution using a Biomek FX with 384-well head (Beckman Coulter, Indianapolis, IN). The DMSO concentration was kept below or at 0.5% v/v. On day 3, rapamycin was added to a final concentration of 30, 0.3, 0.075 (nM), or DMSO control for each screen compound and concentration. To reduce stochastic noise and promote overall screen accuracy, each compound was screened with three concentrations and duplicates.

### Analysis of the screen

Each well of the 384-well plate consists of a population of cells with a specific compound + rapamycin concentration. For each well, we measured the median values for cell size and G1 proportion. We first relied on the rapamycin gradient to produce a linear fit between the size of cells in early G1 (*S*_*EG1*_) and the proportion of cells in early G1 (*P*_*EG1*_). We characterized those that weakened the slope to harbor a *sensor* phenotype. Out of those that maintained a negative slope of *S*_*EG1*_ and *P*_*EG1*_, we further filtered for those that had a significant change in median cell size, which we classify as *dials*.

### siRNA transfection

RPE1-Geminin cells were seeded in 6-well plates at a density of 2 × 10^5^ cells/well. 24 hours post-seeding, the cells were transfected with SMARTpool siRNA (25 nM) for the genes of interest using DharmaFECT one transfection reagent according to the manufacturer's instructions. 20 hours post transfection, the cells were re-suspended using 0.05% trypsin-EDTA (Wisent) and re-seeded into 96-well CellCarrier-96 ultra microplates (PerkinElmer, Woodbridge, ON) at a density of 5,000 cells/well. Six hours after re-seeding, cells were treated with rapamycin (30, 3, 0.3, 0.1 and 0.03 nM) or DMSO control for 24 hr before fixation and staining. The experimental procedures were optimized so there is no observable cell cycle arrest or cell death. Specifically, the siRNA-transfected cells were washed-out of transfection reagent and cultured in regular media when used for the assay with rapamycin concentration series.

### Compound treatment

The cells were seeded into 96-well CellCarrier-96 ultra microplates (PerkinElmer, Woodbridge, ON) 16 hrs prior to drug treatment. To perturb CDK4 activity, Palbociclib and Doxycycline were used at the concentrations indicated in the figure legends. All compounds were dissolved in DMSO and diluted in DMEM when used. The concentrations have been selected to avoid cytostatic effects or cause noticeable cell death. Twenty-four hours post compound treatment, the cells were treated and incubated with rapamycin (30, 3, 0.3, 0.1 and 0.03 nM) for an additional 24 hr before fixation and staining. DMSO only (0.01–0.5% v/v in DMEM) was used as a negative control for all the compound treatment experiments. To examine the process of recovery in cell size, cells were treated with 25nM Torin2 for 24 hr, washed with PBS and re-incubated with conditioned media containing the compounds at the concentrations indicated in the figure legends. For the variation presented in Fig. 3b and Sup Fig. 7a, cells were pre-treated with 25nM Palbociclib or vehicle control for 24 hr prior to Torin2 treatment. Cells were re-suspended using trypsin, and cell size and cell density were measured using the Multisizer 4 Coulter Counter (Beckman-Coulter, Mississauga, ON) or collected for whole cell lysis at time points indicated in the figures. Seeding density was adjusted accordingly to achieve 50-80% confluency at the time of measurement. Vehicle control for all experiments is maintained at 0.01-0.5% v/v DMSO unless otherwise stated.

### Whole cell lysis and western blotting

To prepare whole cell lysates, cells were rinsed with ice-cold PBS and solubilized with RIPA Lysis Buffer (Boston Bio-Products, Boston MA) [50 mM Tris-HCl, 150 mM NaCl, 5 mM EDTA, 1 mM EGTA, 1% NP-40, 0.1% SDS and 0.5% sodium deoxycholate, pH 7.4] supplemented with protease and phosphatase inhibitor Cocktail (Thermo Scientific, Burlington, ON). Protein concentration was determined using the BCA protein assay (Thermo Scientific, Burlington, ON) and suspended with 4X Bolt LDS Sample Buffer and 10X Bolt Reducing Agent and heated for 10 min at 70°C. Samples of equal protein were resolved by SDS-polyacrylamide gel electrophoresis and subjected to immunoblotting for proteins as indicated. The antibodies used for immunoblotting were all purchased from Cell Signaling Technology (Beverly, MA). All Western Blot results in the figures have been reproduced in replicate experiments with cell lysates samples prepared in independent experiments.

### Image processing and cell segmentation

All image analysis (cell segmentation, tracking, measurements of fluorescence intensity and nuclear size) and data analyses were performed with custom-written tools in Matlab. More details can be found in Liu et al.(Liu et al., 2017). To correct for background noise in cell size measurements, we uniformly normalized (subtracted) the minimal SE intensity per unit area.

### Cell cycle stages

Cells were first partitioned, according to their nuclear DNA level, into G1 (2N), S (2N-4N) and G2 (4N) phases. Progression in G1 phase was further divided, based on fluorescence of nuclear Geminin into early G1 (low Geminin), late G1 (medium Geminin), and G1/S transition (higher Geminin). The thresholds were automatically detected based on distribution of DNA and log(Geminin).

For Fig. 2b, measured DAPI intensity values were normalized to center the peaks at 2N and 4N. Since changes in measured DAPI intensity are meaningful representation of cell cycle progression only for cells that are in S phase, but not cells in G1 or G2, we transformed DAPI measurements according to 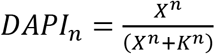 where X are raw measured DAPI intensity values, *K* = 3*N* and *n* = 5. Red circles highlight the size at which 92% of cells have progressed through G1.

### Estimation of cell cycle durations and growth rate from bulk measurements

Cells were treated with inhibitors on multiple 96-well plates and fixed every 20 hr over a period of 3 days. The plate slated to be fixed on the last timepoint were imaged by digital phase contrast (Operetta High-Content Imaging System (PerkinElmer, Woodbridge, ON)) every 12 hrs to acquire cell number estimates. The rate of cell proliferation and protein accumulation were calculated by fitting a log model (assuming that cells proliferate and increase protein content exponentially). More details on growth rate and growth duration calculations can be found in Ginzberg et al.(Ginzberg et al., 2018).

### *C. elegans* maintenance and size measurements

The C. elegans strain JR667 (unc-119(e2498∷Tc1), wIs51[SCMp∷GFP + unc-119(+)]) was provided by the *Caenorhabditis* Genetics Centre, which is funded by NIH Office of Research Infrastructure Programs (P40 OD010440). For RNA interference, JR667 worms were cultivated on lawns of E. coli (strain OP50) expressing double-stranded RNA (dsRNA) from the Source Bioscience LifeSciences feeding library. Individual colonies of E. coli expressing dsRNA were grown to log-phase and platted on nematode growth medium (NGM) plates at 20°C (unless otherwise stated) for 16 h before the addition of the worms. JR667 worms were propagated for two consecutive generations on RNAi bacteria. To measure cell size, second generation worms were synchronized by allowing adult worms to lay eggs for 2 hours, then removing them from the plate. One day after reaching adulthood, second generation worms were mounted on slides in 5μl of 20mM tetramisole anaesthetic on a pad of 4% agar. Z-stack images were taken at 63x using a Zeiss Axio Observer.Z1 microscope. The area of each nucleus/nucleolus was measured at its widest point by hand-tracing the corresponding layer using Image J software.

### Data and Code Availability

Original/source data is available upon request. Further information and requests for resources and reagents should be directed to and will be fulfilled by the lead contact, Ceryl Tan.

## SUPPLEMENTAL INFORMATION TITLES

**Supplementary Figure 1.** Cell size checkpoint in animal cells

**Supplementary Figure 2.** Cell size quantification with SE-A647

**Supplementary Figure 3.** siRNA knockdown of CDK2, CDK4, and CDK6.

**Supplementary Figure 4.** Working concentration of Palbociclib retains active cell cycle.

**Supplementary Figure 5.** Reducing CDK4 activity promotes a rightward shift in the correlation between cell size and G1 length.

**Supplementary Figure 6.** CDK4 activity modulates the proportionality constant for growth rate and growth duration.

**Supplemental Figure 7.** CDK4 activity dials the size threshold at which p38 is activated

**Supplementary Figure 8.** Worm measurements at 1DPA.

**Box 1.** Using single cell measurements to isolate regulatory influences.

## REFERENCES

Allard, C.A.H., Opalko, H.E., Liu, K.-W., Medoh, U., Moseley, J.B., 2018. Cell size-dependent regulation of Wee1 localization by Cdr2 cortical nodes. J. Cell Biol. 217, 1589–1599. https://doi.org/10.1083/jcb.201709171

Altun, Z.F., Hall, D.H., 2002. WormAtlas Hermaphrodite Handbook - Epithelial System - Seam Cells. WormAtlas. https://doi.org/10.3908/wormatlas.1.14

Cadart, C., Monnier, S., Grilli, J., Sáez, P.J., Srivastava, N., Attia, R., Terriac, E., Baum, B., Cosentino-Lagomarsino, M., Piel, M., 2018. Size control in mammalian cells involves modulation of both growth rate and cell cycle duration. Nat. Commun. 9, 3275. https://doi.org/10.1038/s41467-018-05393-0

Canham, S.M., Wang, Y., Cornett, A., Auld, D.S., Baeschlin, D.K., Patoor, M., Skaanderup, P.R., Honda, A., Llamas, L., Wendel, G., Mapa, F.A., Aspesi, P., Labbe-Giguere, N., Gamber, G.G., Palacios, D.S., Schuffenhauer, A., Deng, Z., Nigsch, F., Frederiksen, M., Bushell, S.M., Rothman, D., Jain, R.K., Hemmerle, H., Briner, K., Porter, J.A., Tallarico, J.A., Jenkins, J.L., 2020. Systematic Chemogenetic Library Assembly (preprint). Bioinformatics. https://doi.org/10.1101/2020.03.30.017244

Casanovas, O., Miró, F., Estanyol, J.M., Itarte, E., Agell, N., Bachs, O., 2000. Osmotic stress regulates the stability of cyclin D1 in a p38SAPK2-dependent manner. J. Biol. Chem. 275, 35091–35097. https://doi.org/10.1074/jbc.M006324200

Conlon, I., Raff, M., 2003. Differences in the way a mammalian cell and yeast cells coordinate cell growth and cell-cycle progression. J. Biol. 10.

Cretella, D., Ravelli, A., Fumarola, C., La Monica, S., Digiacomo, G., Cavazzoni, A., Alfieri, R., Biondi, A., Generali, D., Bonelli, M., Petronini, P.G., 2018. The anti-tumor efficacy of CDK4/6 inhibition is enhanced by the combination with PI3K/AKT/mTOR inhibitors through impairment of glucose metabolism in TNBC cells. J. Exp. Clin. Cancer Res. 37, 72. https://doi.org/10.1186/s13046-018-0741-3

Darzynkiewicz, Z., Evenson, D.P., Staiano-Coico, L., Sharpless, T.K., Melamed, M.L., 1979. Correlation between cell cycle duration and RNA content. J. Cell. Physiol. 100, 425–438. https://doi.org/10.1002/jcp.1041000306

Deng, L., Baldissard, S., Kettenbach, A.N., Gerber, S.A., Moseley, J.B., 2014. Dueling kinases regulate cell size at division through the SAD kinase Cdr2. Curr. Biol. CB 24, 428–433. https://doi.org/10.1016/j.cub.2014.01.009

Dolznig, H., Grebien, F., Sauer, T., Beug, H., Müllner, E.W., 2004. Evidence for a size-sensing mechanism in animal cells. Nat. Cell Biol. 6, 899–905. https://doi.org/10.1038/ncb1166

Facchetti, G., Knapp, B., Flor-Parra, I., Chang, F., Howard, M., 2019. Reprogramming Cdr2-Dependent Geometry-Based Cell Size Control in Fission Yeast. Curr. Biol. CB 29, 350–358.e4. https://doi.org/10.1016/j.cub.2018.12.017

Flemming, A.J., Shen, Z.Z., Cunha, A., Emmons, S.W., Leroi, A.M., 2000. Somatic polyploidization and cellular proliferation drive body size evolution in nematodes. Proc. Natl. Acad. Sci. U. S. A. 97, 5285–5290. https://doi.org/10.1073/pnas.97.10.5285

Franco, J., Balaji, U., Freinkman, E., Witkiewicz, A.K., Knudsen, E.S., 2016. Metabolic Reprogramming of Pancreatic Cancer Mediated by CDK4/6 Inhibition Elicits Unique Vulnerabilities. Cell Rep. 14, 979–990. https://doi.org/10.1016/j.celrep.2015.12.094

Gao, F.B., Raff, M., 1997. Cell size control and a cell-intrinsic maturation program in proliferating oligodendrocyte precursor cells. J. Cell Biol. 138, 1367–1377. https://doi.org/10.1083/jcb.138.6.1367

Gerganova, V., Floderer, C., Archetti, A., Michon, L., Carlini, L., Reichler, T., Manley, S., Martin, S.G., 2019. Multi-phosphorylation reaction and clustering tune Pom1 gradient mid-cell levels according to cell size. eLife 8. https://doi.org/10.7554/eLife.45983

Ginzberg, M.B., Chang, N., D’Souza, H., Patel, N., Kafri, R., Kirschner, M.W., 2018. Cell size sensing in animal cells coordinates anabolic growth rates and cell cycle progression to maintain cell size uniformity. eLife 7, e26957. https://doi.org/10.7554/eLife.26957

Haines, E., Chen, T., Kommajosyula, N., Chen, Z., Herter-Sprie, G.S., Cornell, L., Wong, K.-K., Shapiro, G.I., 2018. Palbociclib resistance confers dependence on an FGFR-MAP kinase-mTOR-driven pathway in *KRAS* -mutant non-small cell lung cancer. Oncotarget 9, 31572–31589. https://doi.org/10.18632/oncotarget.25803

Hartwell, L.H., Culotti, J., Pringle, J.R., Reid, B.J., 1974. Genetic control of the cell division cycle in yeast. Science 183, 46–51.

Helman, A., Klochendler, A., Azazmeh, N., Gabai, Y., Horwitz, E., Anzi, S., Swisa, A., Condiotti, R., Granit, R.Z., Nevo, Y., Fixler, Y., Shreibman, D., Zamir, A., Tornovsky-Babeay, S., Dai, C., Glaser, B., Powers, A.C., Shapiro, A.M.J., Magnuson, M.A., Dor, Y., Ben-Porath, I., 2016. p16(Ink4a)-induced senescence of pancreatic beta cells enhances insulin secretion. Nat. Med. 22, 412–420. https://doi.org/10.1038/nm.4054

Jansen, V.M., Bhola, N.E., Bauer, J.A., Formisano, L., Lee, K.-M., Hutchinson, K.E., Witkiewicz, A.K., Moore, P.D., Estrada, M.V., Sánchez, V., Ericsson, P.G., Sanders, M.E., Pohlmann, P.R., Pishvaian, M.J., Riddle, D.A., Dugger, T.C., Wei, W., Knudsen, E.S., Arteaga, C.L., 2017. Kinome-Wide RNA Interference Screen Reveals a Role for PDK1 in Acquired Resistance to CDK4/6 Inhibition in ER-Positive Breast Cancer. Cancer Res. 77, 2488–2499. https://doi.org/10.1158/0008-5472.CAN-16-2653

Kafri, R., Levy, J., Ginzberg, M.B., Oh, S., Lahav, G., Kirschner, M.W., 2013. Dynamics extracted from fixed cells reveal feedback linking cell growth to cell cycle. Nature 494, 480–483. https://doi.org/10.1038/nature11897

Killander, D., Zetterberg, A., 1965. QUANTITATIVE CYTOCHEMICAL STUDIES ON INTERPHASE GROWTH. DETERMINATION OF DNA, RNA AND MASS CONTENT OF AGE DETERMINED MOUSE FIBROBLASTS IN VITRO AND OF INTERCELLULAR VARIATION IN GENERATION TIME. Exp. Cell Res. 38, 272–284. https://doi.org/10.1016/0014-4827(65)90403-9

Lee, Y., Dominy, J.E., Choi, Y.J., Jurczak, M., Tolliday, N., Camporez, J.P., Chim, H., Lim, J.-H., Ruan, H.-B., Yang, X., Vazquez, F., Sicinski, P., Shulman, G.I., Puigserver, P., 2014. Cyclin D1–Cdk4 controls glucose metabolism independently of cell cycle progression. Nature 510, 547–551. https://doi.org/10.1038/nature13267

Leitao, R.M., Kellogg, D.R., 2017. The duration of mitosis and daughter cell size are modulated by nutrients in budding yeast. J. Cell Biol. 216, 3463–3470. https://doi.org/10.1083/jcb.201609114

Liu, G.Y., Sabatini, D.M., 2020. mTOR at the nexus of nutrition, growth, ageing and disease. Nat. Rev. Mol. Cell Biol. 21, 183–203. https://doi.org/10.1038/s41580-019-0199-y

Liu, S., Ginzberg, M.B., Patel, N., Hild, M., Leung, B., Li, Z., Chen, Y.-C., Chang, N., Wang, Y., Tan, C., Diena, S., Trimble, W., Wasserman, L., Jenkins, J.L., Kirschner, M.W., Kafri, R., 2018. Size uniformity of animal cells is actively maintained by a p38 MAPK-dependent regulation of G1-length. eLife 7, e26947. https://doi.org/10.7554/eLife.26947

Llanos, S., Megias, D., Blanco-Aparicio, C., Hernández-Encinas, E., Rovira, M., Pietrocola, F., Serrano, M., 2019. Lysosomal trapping of palbociclib and its functional implications. Oncogene 38, 3886–3902. https://doi.org/10.1038/s41388-019-0695-8

Lloyd, A.C., 2013. The Regulation of Cell Size. Cell 154, 1194–1205. https://doi.org/10.1016/j.cell.2013.08.053

Lopez-Mejia, I.C., Lagarrigue, S., Giralt, A., Martinez-Carreres, L., Zanou, N., Denechaud, P.-D., Castillo-Armengol, J., Chavey, C., Orpinell, M., Delacuisine, B., Nasrallah, A., Collodet, C., Zhang, L., Viollet, B., Hardie, D.G., Fajas, L., 2017. CDK4 Phosphorylates AMPKα2 to Inhibit Its Activity and Repress Fatty Acid Oxidation. Mol. Cell 68, 336–349.e6. https://doi.org/10.1016/j.molcel.2017.09.034

Malumbres, M., Barbacid, M., 2009. Cell cycle, CDKs and cancer: a changing paradigm. Nat. Rev. Cancer 9, 153–166. https://doi.org/10.1038/nrc2602

Martin, S.G., Berthelot-Grosjean, M., 2009. Polar gradients of the DYRK-family kinase Pom1 couple cell length with the cell cycle. Nature 459, 852–856. https://doi.org/10.1038/nature08054

Meyer, C.A., 2000. Drosophila Cdk4 is required for normal growth and is dispensable for cell cycle progression. EMBO J. 19, 4533–4542. https://doi.org/10.1093/emboj/19.17.4533

Moseley, J.B., Mayeux, A., Paoletti, A., Nurse, P., 2009. A spatial gradient coordinates cell size and mitotic entry in fission yeast. Nature 459, 857–860. https://doi.org/10.1038/nature08074

Neufeld, T.P., de la Cruz, A.F., Johnston, L.A., Edgar, B.A., 1998. Coordination of growth and cell division in the Drosophila wing. Cell 93, 1183–1193. https://doi.org/10.1016/s0092-8674(00)81462-2

Nurse, P., 1975. Genetic control of cell size at cell division in yeast. Nature 256, 547–551. https://doi.org/10.1038/256547a0

Park, M., Krause, M.W., 1999. Regulation of postembryonic G(1) cell cycle progression in Caenorhabditis elegans by a cyclin D/CDK-like complex. Dev. Camb. Engl. 126, 4849–4860.

Regot, S., Hughey, J.J., Bajar, B.T., Carrasco, S., Covert, M.W., 2014. High-sensitivity measurements of multiple kinase activities in live single cells. Cell 157, 1724–1734. https://doi.org/10.1016/j.cell.2014.04.039

Romero-Pozuelo, J., Figlia, G., Kaya, O., Martin-Villalba, A., Teleman, A.A., 2020. Cdk4 and Cdk6 Couple the Cell-Cycle Machinery to Cell Growth via mTORC1. Cell Rep. 31, 107504. https://doi.org/10.1016/j.celrep.2020.03.068

Schaechter, M., Williamson, J.P., Hood, J.R., Koch, A.L., 1962. Growth, cell and nuclear divisions in some bacteria. J. Gen. Microbiol. 29, 421–434. https://doi.org/10.1099/00221287-29-3-421

Schmoller, K.M., Turner, J.J., Kõivomägi, M., Skotheim, J.M., 2015. Dilution of the cell cycle inhibitor Whi5 controls budding-yeast cell size. Nature 526, 268–272. https://doi.org/10.1038/nature14908

Sellam, A., Chaillot, J., Mallick, J., Tebbji, F., Richard Albert, J., Cook, M.A., Tyers, M., 2019. The p38/HOG stress-activated protein kinase network couples growth to division in Candida albicans. PLOS Genet. 15, e1008052. https://doi.org/10.1371/journal.pgen.1008052

Shields, R., Brooks, R.F., Riddle, P.N., Capellaro, D.F., Delia, D., 1978. Cell size, cell cycle and transition probability in mouse fibroblasts. Cell 15, 469–474. https://doi.org/10.1016/0092-8674(78)90016-8

Thoms, H.C., Dunlop, M.G., Stark, L.A., 2007. p38-mediated inactivation of cyclin D1/cyclin-dependent kinase 4 stimulates nucleolar translocation of RelA and apoptosis in colorectal cancer cells. Cancer Res. 67, 1660–1669. https://doi.org/10.1158/0008-5472.CAN-06-1038

Tzur, A., Kafri, R., LeBleu, V.S., Lahav, G., Kirschner, M.W., 2009. Cell growth and size homeostasis in proliferating animal cells. Science 325, 167–171. https://doi.org/10.1126/science.1174294

Varsano, G., Wang, Y., Wu, M., 2017. Probing Mammalian Cell Size Homeostasis by Channel-Assisted Cell Reshaping. Cell Rep. 20, 397–410. https://doi.org/10.1016/j.celrep.2017.06.057

Wang, C., Li, Z., Lu, Y., Du, R., Katiyar, S., Yang, J., Fu, M., Leader, J.E., Quong, A., Novikoff, P.M., Pestell, R.G., 2006. Cyclin D1 repression of nuclear respiratory factor 1 integrates nuclear DNA synthesis and mitochondrial function. Proc. Natl. Acad. Sci. U. S. A. 103, 11567–11572. https://doi.org/10.1073/pnas.0603363103

Weber, S.C., Brangwynne, C.P., 2015. Inverse Size Scaling of the Nucleolus by a Concentration-Dependent Phase Transition. Curr. Biol. 25, 641–646. https://doi.org/10.1016/j.cub.2015.01.012

Willis, L., Refahi, Y., Wightman, R., Landrein, B., Teles, J., Huang, K.C., Meyerowitz, E.M., Jönsson, H., 2016. Cell size and growth regulation in the *Arabidopsis thaliana* apical stem cell niche. Proc. Natl. Acad. Sci. 113, E8238–E8246. https://doi.org/10.1073/pnas.1616768113

Wood, E., Nurse, P., 2013. Pom1 and cell size homeostasis in fission yeast. Cell Cycle 12, 3417–3425. https://doi.org/10.4161/cc.26462

Xie, S., Skotheim, J.M., 2020. A G1 Sizer Coordinates Growth and Division in the Mouse Epidermis. Curr. Biol. 30, 916–924.e2. https://doi.org/10.1016/j.cub.2019.12.062

Zatulovskiy, E., Skotheim, J.M., 2020. On the Molecular Mechanisms Regulating Animal Cell Size Homeostasis. Trends Genet. 36, 360–372. https://doi.org/10.1016/j.tig.2020.01.011

Zatulovskiy, E., Zhang, S., Berenson, D.F., Topacio, B.R., Skotheim, J.M., 2020. Cell growth dilutes the cell cycle inhibitor Rb to trigger cell division 6.

Zhang, C., Stockwell, S.R., Elbanna, M., Ketteler, R., Freeman, J., Al-Lazikani, B., Eccles, S., De Haven Brandon, A., Raynaud, F., Hayes, A., Clarke, P.A., Workman, P., Mittnacht, S., 2019. Signalling involving MET and FAK supports cell division independent of the activity of the cell cycle-regulating CDK4/6 kinases. Oncogene 38, 5905–5920. https://doi.org/10.1038/s41388-019-0850-2

