## Supplemental Figures for "Cell size homeostasis is maintained by a circuitry involving a CDK4-determined target size that programs the cell size-dependent activation of p38"

### SUPPLEMENTAL INFORMATION

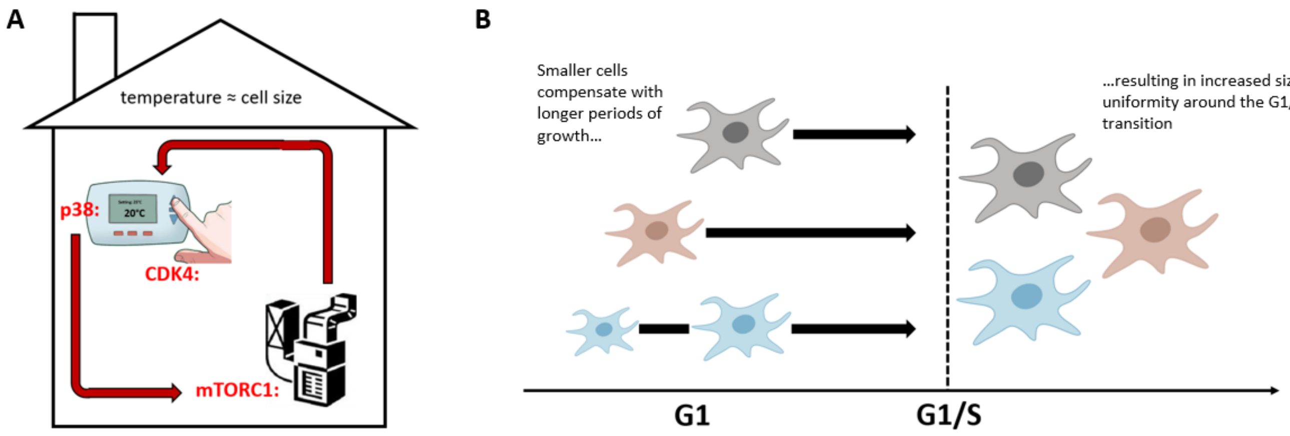


**Supplementary Figure 1. Cell size checkpoint in animal cells (A)** Proposed model for animal cell size checkpoint where cell size homeostasis is maintained similarly to the function of a thermostat – p38 is analogous to the sensor (or thermometer), mTORC1 to the furnace, and CDK4 to the thermostat dial that sets the critical target size. **(B)** Cells that were born smaller tend to have a longer G1 phase, resulting in increased uniformity in the cells undergoing G1/S transition.

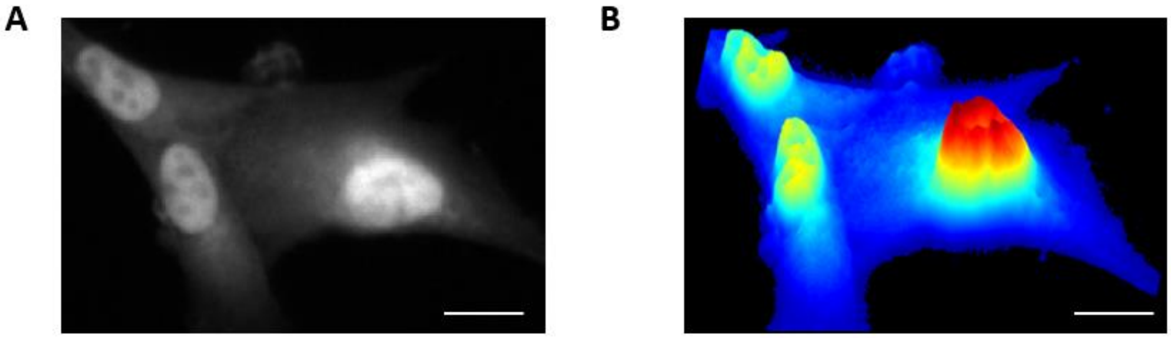


**Supplementary Figure 2. Cell size quantification with SE-A647 (A)** Widefield fluorescence image of RPE1 cells stained with succinimidyl ester conjugated with an Alexa Fluor 647 (SE-A647) stain. **(B)** 3D projection of the same cells, with the fluorescence intensity of SE-A647 depicted in blue to red for low to high. Scale bar = 12.5μm.

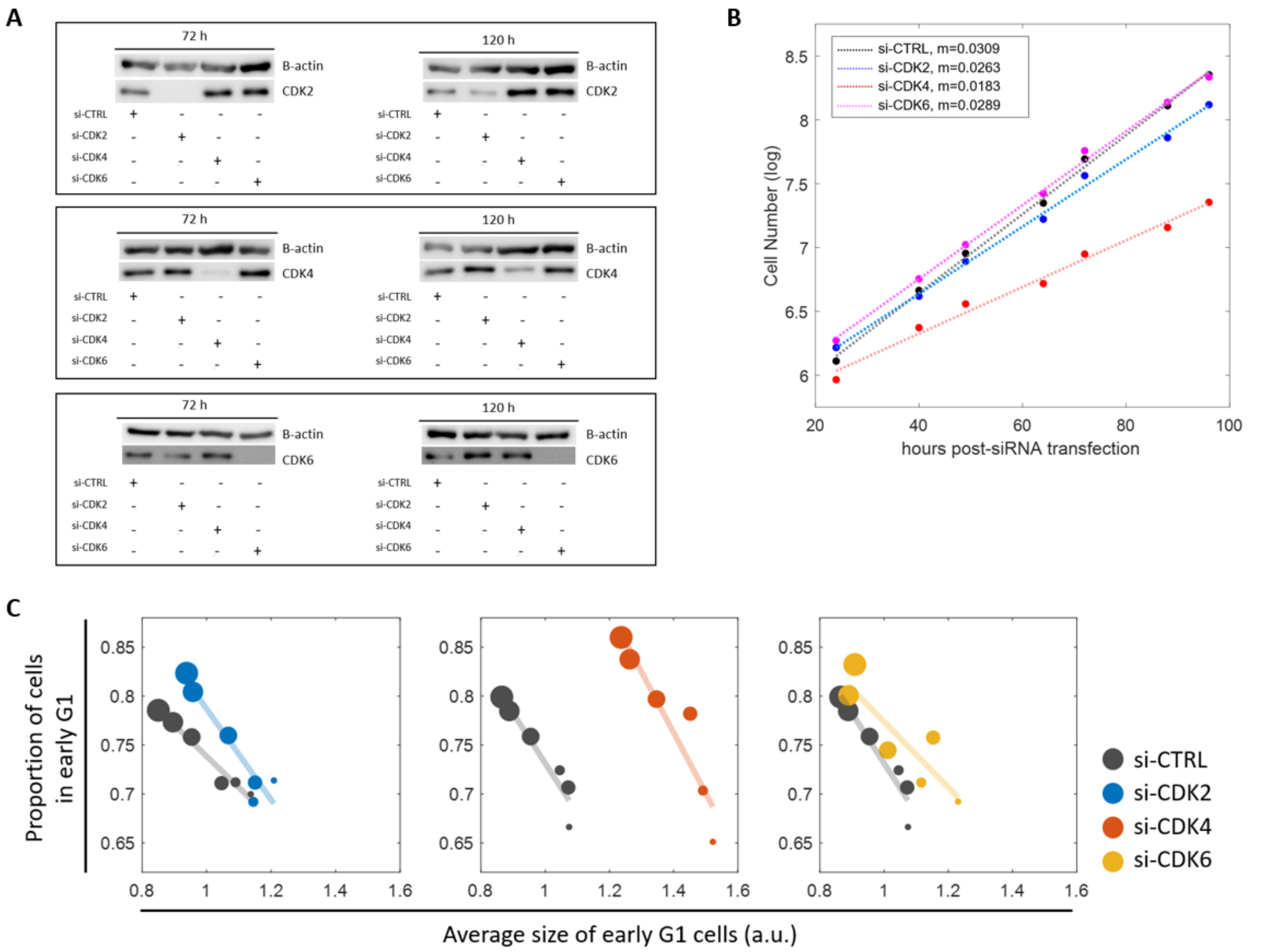


**Supplementary Figure 3. siRNA knockdown of CDK2, CDK4, and CDK6. (A)** Western blot validation of siRNA against CDK2, CDK4, and CDK6 at 72 hrs or 120 hrs post-transfection. B-actin is shown as loading control. **(B)** Active proliferation is shown as a linear fit between cell number (in log scale) and time, with the slope (m) showing the rate of proliferation. **(C)** Knockdown of all 3 CDKs retain a negative correlation between cell size and G1 length, but only si-CDK4 shifts the coordination. Proportion of cells in G1 is used as a proxy for G1 length.

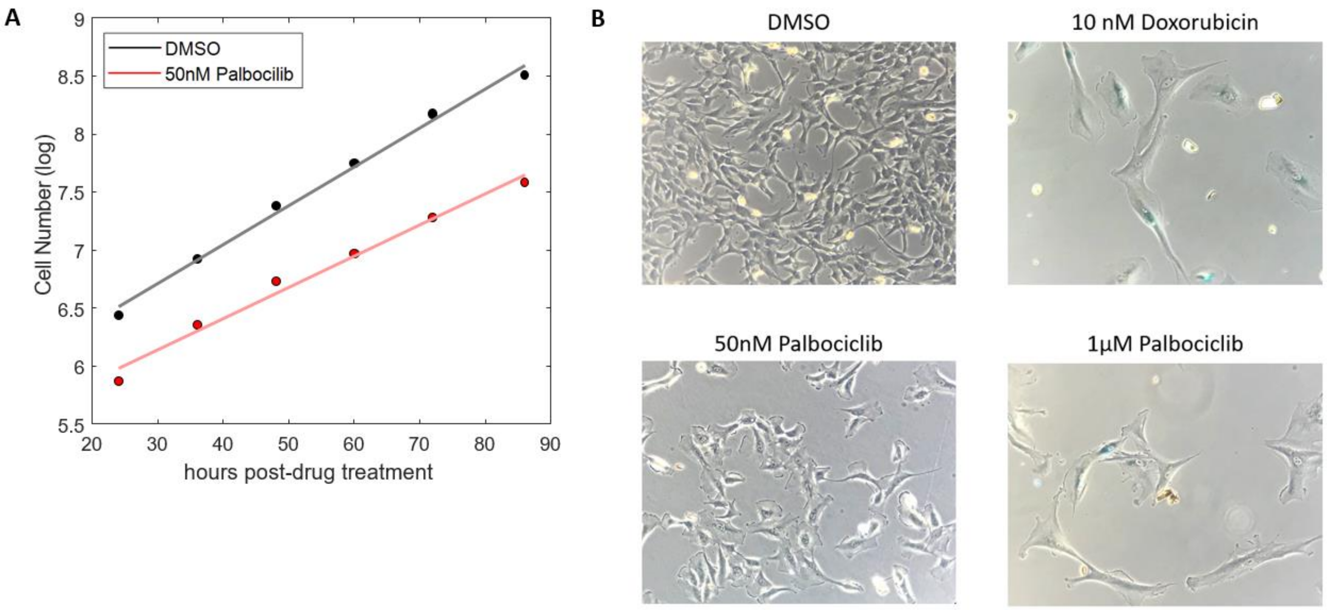


**Supplementary Figure 4. Working concentration of Palbociclib retains active cell cycle. (A)** Cells treated with 50nM Palbociclib maintain exponential proliferation (at a slower rate than control) as shown by a linear fit of cell number (in log scale) with time. **(B)** Beta-galactosidase staining on RPE1 cells incubated with the various conditions for 7 days. Blue staining (indicative of senescence) can be observed with 1µM Palbociclib and 10nM doxorubicin, but not 50nm Palbociclib. All wells were seeded at the same density.

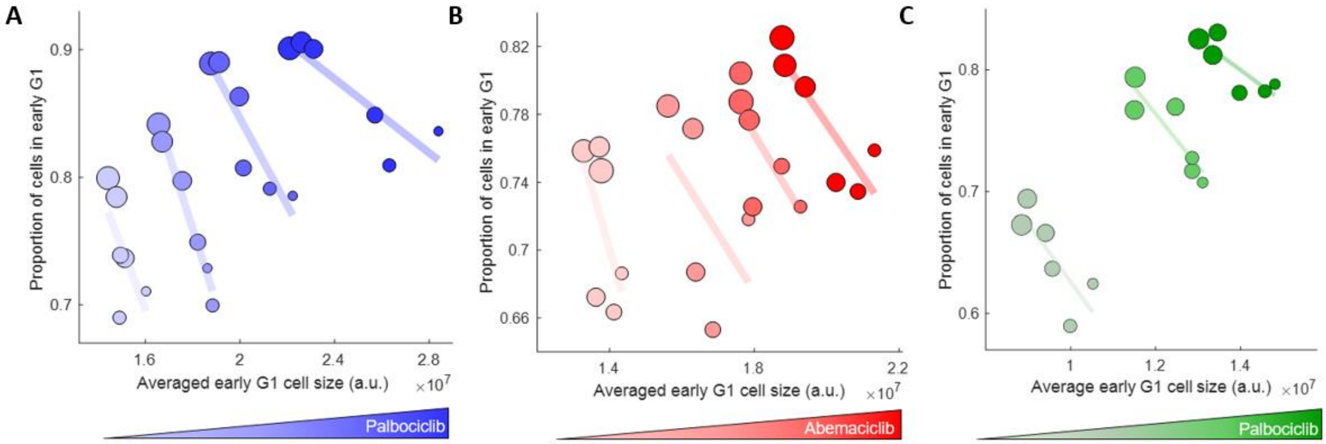


**Supplementary Figure 5. Reducing CDK4 activity promotes a rightward shift in the correlation between cell size and G1 length.** Increasing concentrations of **(A)** palbociclib (0, 12.5nM, 25nM, 50nM) and **(B)** abemaciclib (0, 10nM, 25nM, 50nM), both of which are selective inhibitors of CDK4/6, show the same trend. **(C)** Primary human fibroblasts treated with 0, 50nM and 100nM palbociclib also show the same trend. Proportion of cells in G1 is used as a proxy for G1 length.

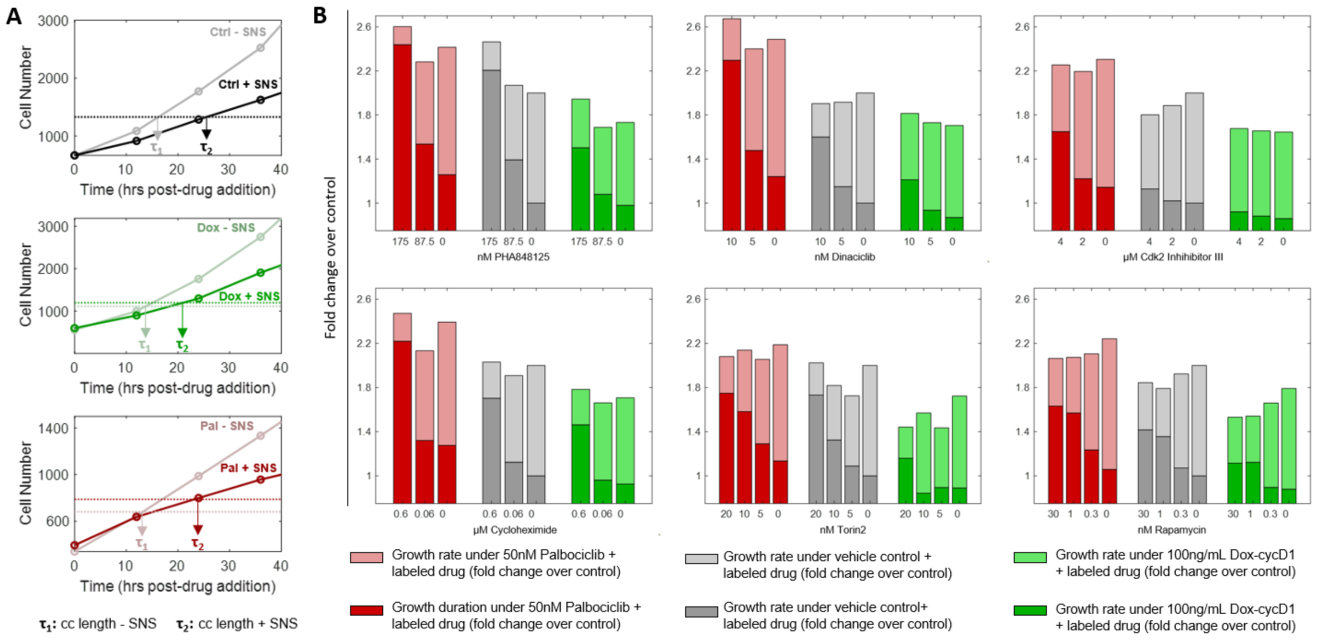


**Supplementary Figure 6. CDK4 activity modulates the proportionality constant for growth rate and growth duration. (A)** Proliferation curves for the different combinations from Fig. 2D, with dotted lines depicting doubling time for each background level of CDK4 activity (τ1) and under co-treatment with 39nM SNS-032 (τ2). **(B)** Changes in growth rate (lighter bars) and cell cycle length (darker bars) presented as fold change to control for the different backgrounds under various perturbations on growth duration (top) or growth rate (bottom). Height of bars represents the cell size achieved (control=2).

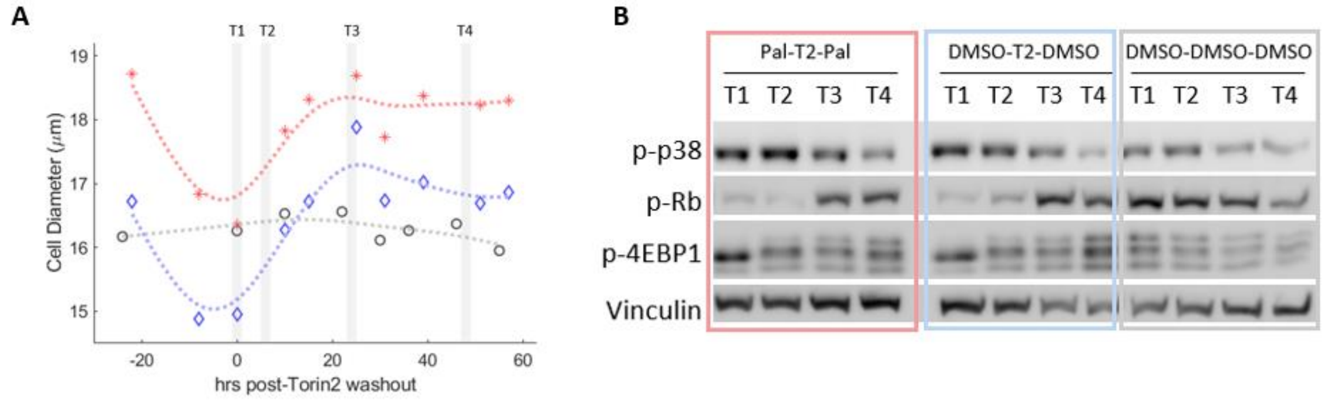


**Supplemental Figure 7. CDK4 activity dials the size threshold at which p38 is activated (A)** RPE1 cells were pre-treated with 25nM Palbociclib or vehicle control for 24hr prior to the Torin2-recovery assay, and p38 activity was measured from lysates collected at 4 timepoints during the recovery phase. **(B)** Western blots of lysates are shown.

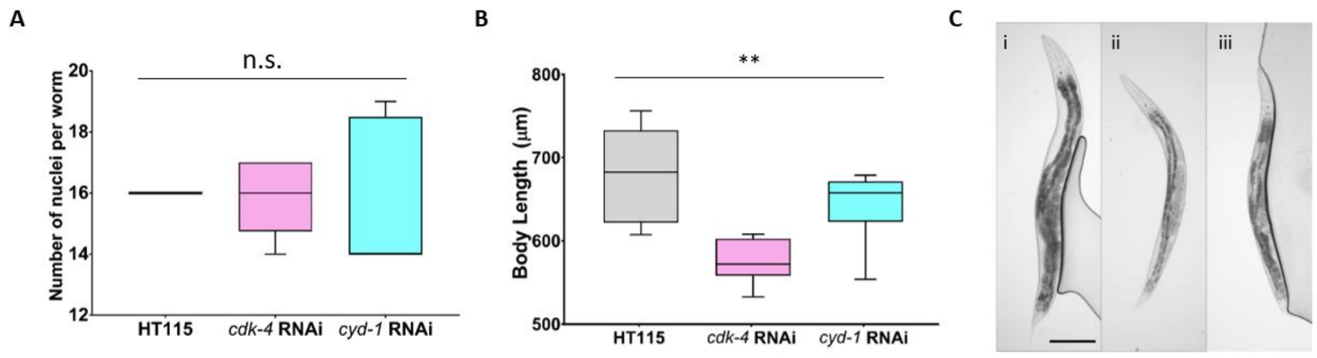


**Supplementary Figure 8. Worm measurements at 1DPA. (A)** Number of seam cell nuclei along one side per worm (one nucleolus per nucleus). **(B)** Body length measurements of JR667 worms on RNAi. Target gene knockdown decreased body length compared to control worms of the same life stage. **(C)** Representative images of JR667 worms at the one-day post-adult stage (1DPA) on different RNAi treatments. Images depict (i) HT115 control, (ii) *cdk-4* RNAi and (iii) *cyd-1* RNAi. Scale bar = 100µm. HT115 RNAi n=6 worms, *cdk-4* RNAi n = 9, *cyd-1* RNAi n = 5. ***p<0.001, unpaired t-test.

**Box 1**

**Using single cell measurements to isolate regulatory influences.** A hypothesis investigated in the present manuscript states that: (A) p38MAPK is activated in a cell size dependent manner, and that (B) the mutual dependency of p38 activity and cell size is 'programmed' by CDK4. Nonetheless, it would be wrong to assume that size regulation is the sole function of p38. The question, therefore, is how to measure the mutual dependency of p38 and cell size, while normalizing for other regulatory influences that affect these processes? Similarly, with chemical or genetic perturbations, our question is not how these perturbations affect p38 activity or cell size but rather – to measure how these perturbations affect the joint dependency of size and p38. To address these challenges, we use single cell data to compare **two** separate probability distributions:

| | Probability distribution | Description |
|---|---|---|
| *1* | $f_{joint} = f(p38KTR, size)$ | The measured joint distribution quantifies the frequency of cells with any given paired cell size and p38-KTR values. |
| *2* | $f_{ind} = f(p38KTR) \times f(size)$ | The product of the measured distribution (marginal) of p38 KTR and the measured (marginal) distribution of cell size. A fundamental principle in statistics states that, if two variables (in our case, p38-KTR and cell size) are independent, their joint probability distribution is precisely the product of the marginal distributions. Therefore, $f_{ind}$ precisely represents what the joint distribution ($f_{joint}$) of p38-KTR and cell size ***would*** have looked like, had the two been ***independent***. Both $f_{joint}$ and $f_{ind}$ are empirically measured from single cell measurements. This is important because, the differences between $f_{joint}$ and $f_{ind}$ effectively quantifies the mutual dependency of p38 activity and cell size – presumably a dependency that is established by intracellular regulation linking the two processes |

With the two measured distributions above, we perform two comparisons: the first characterizing the mutual dependency of p38-KTR and cell size, while the second asks how this mutual dependency is influenced by chemical or genetic perturbations.

To quantify the mutual dependency of cell size and p38-KTR, we normalize the measured distribution, $f_{joint}$, to what that distribution would have looked like had p38 and size been independent ($f_{ind}$). As a measure of comparison, we use the log ratio to define $\delta_{reg}$:

$$\delta_{reg} = log\left(\frac{f_{joint}}{f_{ind}}\right) = log\left(\frac{f(p38KTR, size)}{f(p38KTR) \times f(size)}\right)$$

Essentially, $\delta_{reg}$ is a heatmap that quantifies whether cells with any given size and p38 activity levels are over- or under- represented as compared to the $f_{ind}$. To emphasize, since both $f_{ind}$ and $f_{joint}$ are empirically measured, $\delta_{reg}$ is an empirically measured quantity that describes the mutual dependency of size and p38. In other words, $\delta_{reg}$ is a measurement of the regulatory dependency between size and p38 – a dependency that is assumed to reflect an intracellular regulation linking the two.

To measure how drug treatments affect the mutual dependency of size and p38 activity levels, we calculate $\delta_{reg}$ from measurements on cells subject to drug treatments. Figure 3f highlights the utility of $\delta_{reg}$ as a measure of regulatory interactions. While rapamycin affects cell size, rapamycin does not affect the measured $\delta_{reg}$ values. Because $\delta_{reg}$ is normalized to both $f(p38KTR)$ and $f(size)$, $\delta_{reg}$ values will not change by perturbations that change cell size and/or p38, so long as these perturbations do not influence the mutual dependence of size and p38.
